# High-throughput generic single-entity sequencing using droplet microfluidics

**DOI:** 10.1101/2023.08.15.549386

**Authors:** Guoping Wang, Liuyang Zhao, Yu Shi, Fuyang Qu, Yanqiang Ding, Weixin Liu, Changan Liu, Gang Luo, Meiyi Li, Xiaowu Bai, Luoquan Li, Yi-Ping Ho, Jun Yu

**Affiliations:** Institute of Digestive Diseases and Department of Medicine and Therapeutics, State Key Laboratory of Digestive Disease, Li Ka Shing Institute of Health Sciences, The Chinese University of Hong Kong, Hong Kong SAR, China; Department of Biomedical Engineering, The Chinese University of Hong Kong, Hong Kong SAR, China; Department of Electronic Engineering, The Chinese University of Hong Kong, Hong Kong SAR, China; Hong Kong Branch of CAS Center for Excellence in Animal Evolution and Genetics, The Chinese University of Hong Kong, Hong Kong SAR, China; The Ministry of Education Key Laboratory of Regeneration Medicine, The Chinese University of Hong Kong, Hong Kong SAR, China

## Abstract

Single-cell sequencing has revolutionized our understanding of cellular heterogeneity by providing a micro-level perspective over the past decades. Although heterogeneity is essential for various biological communities, the currently demonstrated platform predominantly focuses on eukaryotic cells without cell walls and their transcriptomics ^1,2^, leaving significant gaps in the study of omics from other single biological entities such as bacteria and viruses. Due to the difficulty of isolating and acquiring their DNA^3^, contemporary methodologies for the characterization of generic biological entities remain conspicuously constrained, with low throughput^4^, compromised lysis efficiency^5^, and highly fragmented genomes^6^. Herein, we present the Generic Single Entity Sequencing platform (GSE-Seq), which boasts ample versatility, high throughput, and high coverage, and is enabled by an innovative workflow, addressing the critical challenges in single entities sequencing: (1) one-step manufacturing of massive barcodes, (2) degradable hydrogel-based *in situ* sample processing and whole genome amplification, (3) integrated in-drop library preparation, (4) compatible long-read sequencing. By GSE-Seq, we have achieved a significant milestone by enabling high-throughput, long-read single-entity profiling of dsDNA and ssDNA from single virus sequencing (SV-seq) and single bacteria sequencing (SB-seq) of the human gut and marine sediment for the first time. Notably, our analysis uncovered previously overlooked viral and bacterial dark matter and phage-host interactions. In summary, the presented conceptually new workflow offers a toolbox based on droplet microfluidics to tackle the persistent challenges in high-throughput profiling to generic applications, which hold immense promise for diverse biological entities, especially hard-to-lyse cells.

## Main

The biological realm encompasses an extremely large number and diverse array of entities, including eukaryotic cells, prokaryotic cells, and virus particles. Obtaining the complete genome in a high-throughput manner is crucial for comprehensively understanding their biology. Genomes can comprise different types of DNA, including double-stranded (dsDNA), single-stranded (ssDNA), linear, and circular DNA. Among these entities, analyzing a single microbiome is notably challenging due to the significantly smaller size of bacterial (about 1000 times) and viral genomes (around 10000 times) compared to human cells. Additionally, the complex cell structure of microbiomes further complicates lysis.

While droplet microfluidics featured in high-throughput analysis has shown significant promise for single-cell sequencing, challenges remain when multi-step processes are involved, owing to the instability of water-oil interface susceptible to the harsh lysis conditions, multiple-cycle enzyme treatments, and purification steps associated with the processing of samples. As a result, sequencing a single microbiome and other hard-to-lyse cells lags behind mammalian cells. To address these challenges and expand single-cell analysis to a broader range of biological entities, we present a novel Generic Single Entity Sequencing (GSE-Seq) platform for massive genome sequencing of individual entities.

GSE-Seq, featured in ample versatility, high throughput, and high coverage, is enabled by an innovatively designed workflow, addressing the critical challenges in single entities sequencing: (1) degradable hydrogel-based *in situ* sample processing and whole genome amplification, (2) one-step manufacturing of massive barcode, (3) integrated in-drop library preparation, (4) compatible long-read sequencing (Figure 1a). Enabled by the seamlessly integrated workflow and well-optimized microfluidic modules, GSE-Seq is promising for single biological entity sequencing, particularly for long-read sequencing.

**Figure 1.**
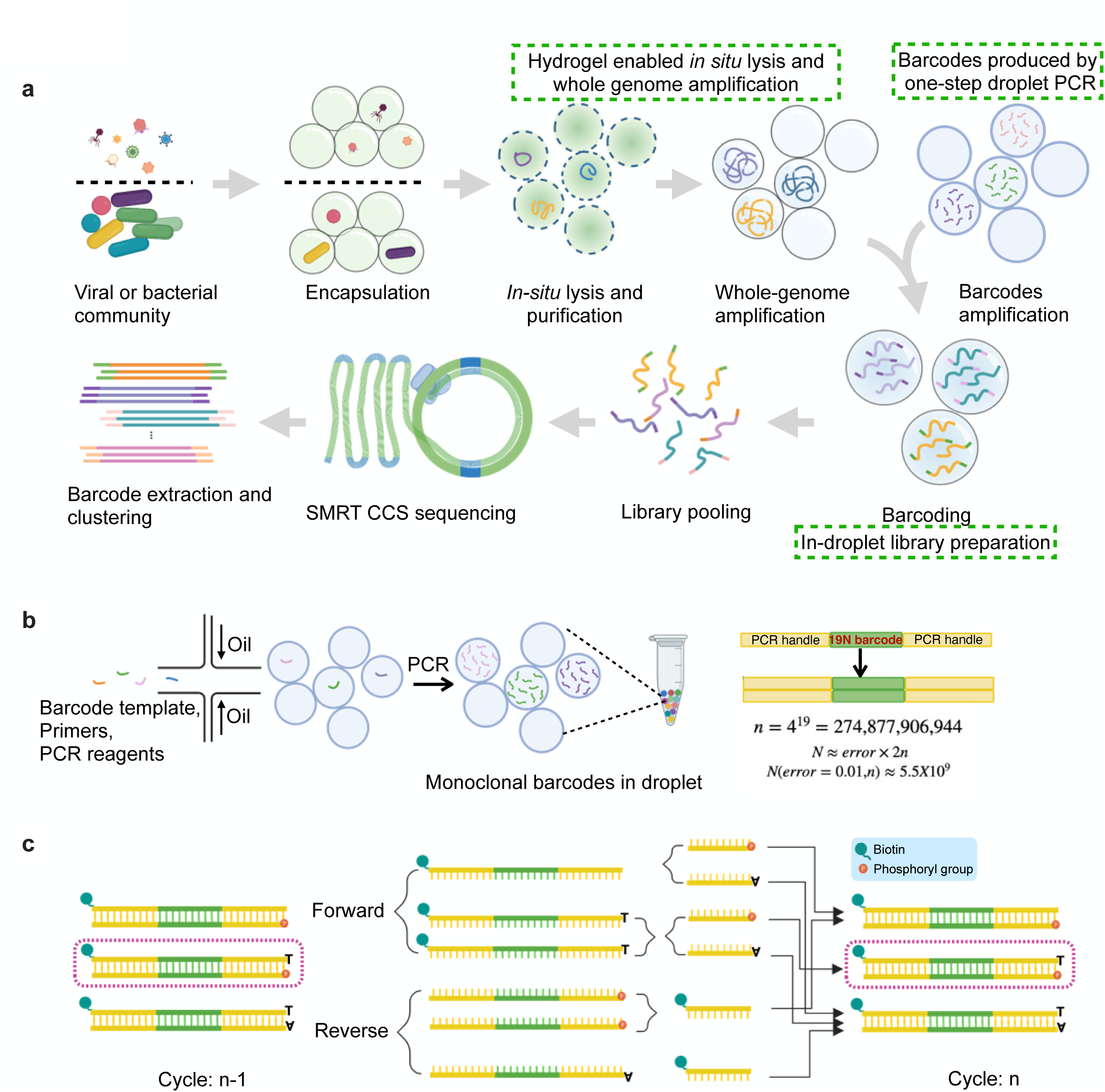
A fast, low-cost, high-throughput barcode generation for massive single-entity sequencing. **a,** Schematic of the GSE-seq workflow. **b,** Barcode pool generated by one-step droplet PCR. **c,** Barcode generation in each PCR cycle.

### Rapid, low-cost, and versatile generation of massive monoclonal barcodes by one-step droplet PCR

The key principle of high-throughput single-cell genomics analysis hinges on high-efficiency barcoding, which involves labelling the nucleic acid originating in the same genome with a unique identifier. Classic barcoding strategies employ split-and-pool oligo synthesis on hydrogel beads, which is labor-intensive, costly, and inefficient, posing difficulties for implementation in many laboratories^7^. To address this challenge, a brand-new droplet barcode was developed as a convenient and low-cost approach for generating more than billions of barcode droplets, each containing a distinct clonal population of a barcode (Figure 1b, Video S1). Within each droplet, the PCR reagents and barcode template were mixed with specially designed three primers —a forward Primer A with a biotinylated 5’-terminal, two reverse primers, Primer B with 5’-terminal phosphorylation, and Primer C with an extra dA at the 5’-terminal compared to the Primer B (Figure S1a, Table S1). Upon in-droplet PCR amplification, the barcode oligo – a sequence of random bases flanked by two PCR handles – can generate an attachable barcode, which is biotinylated, double-stranded, and has a phosphorylated end with a protruding dT (Figure 1b,c, S1a-c). Nineteen random bases were designed to theoretically produce about 2.7×10^11^ barcode combinations, generating a barcode pool of more than 5×10^9^ with a tolerance error rate^8^ of less than 1% (Figure 1b). The proposed design significantly increases the diversity of the barcode pool.

Compared to previous *in-situ* synthesis barcodes on beads, barcodes in droplets, as demonstrated here, make synthesis and purification easier, resulting in a lower error rate^9^, which is particularly important for long DNA barcodes. We synthesized our barcode oligos with a commonly used DNA synthesizer or through commercially available service in less than 100 dollars (Table S1). In the meantime, one can tailored-modify the barcode design facilely to suit various applications. With simple one-step in-droplet PCR at a low cost, billions of barcodes can be generated within hours. Furthermore, the rapid generation of sticky ends is also made possible by large-scale DNA cloning, which promotes this entirely new barcode strategy for generic applications.

### Novel degradable hydrogel beads enabled *in situ* lysis and highly efficient whole genome amplification

Water-in-oil droplets made by surfactants typically used for single-cell sequencing are susceptible to breakage, particularly when exposed to certain chemicals targeting hard-to-lyse entities such as bacteria that necessitate harsh lysis conditions, including higher detergent concentrations. In a previous study, porous agarose matrices were employed to encapsulate bacteria for subsequent lysis^6^. However, the degradation of the agarose matrix posed significant obstacles for downstream reactions, such as whole genome amplification (WGA). To this end, GSE-Seq specifically incorporated a degradable hydrogel that is not only compatible with harsh lysis conditions and multiple-step DNA purification but also easy to degrade, allowing GSE-Seq generally applicable to a variety of downstream reactions.

Massive isolation of single entities, *in situ* lysis, genome purification, and WGA were demonstrated using degradable hydrogel coupled with droplet microfluidics (Figure 2a). Enzyme-mediated crosslink chemistry^10^ was employed to simplify the operational protocol and minimize the risk of DNA damage (Figure 2a, S2a). By taking advantage of this approach, the gelation did not necessitate using specialized heating equipment typically required for agarose or others^6^. The prepared aqueous hydrogel solution can be simply cooled using an ice bag to slow down the crosslinking reaction during droplet generation to enable the massive encapsulation of single entities. Upon encapsulation, the produced droplets (>100 million) containing single entities were incubated at room temperature for the gelation (Figure 2a, b, Video S2).

**Figure 2.**
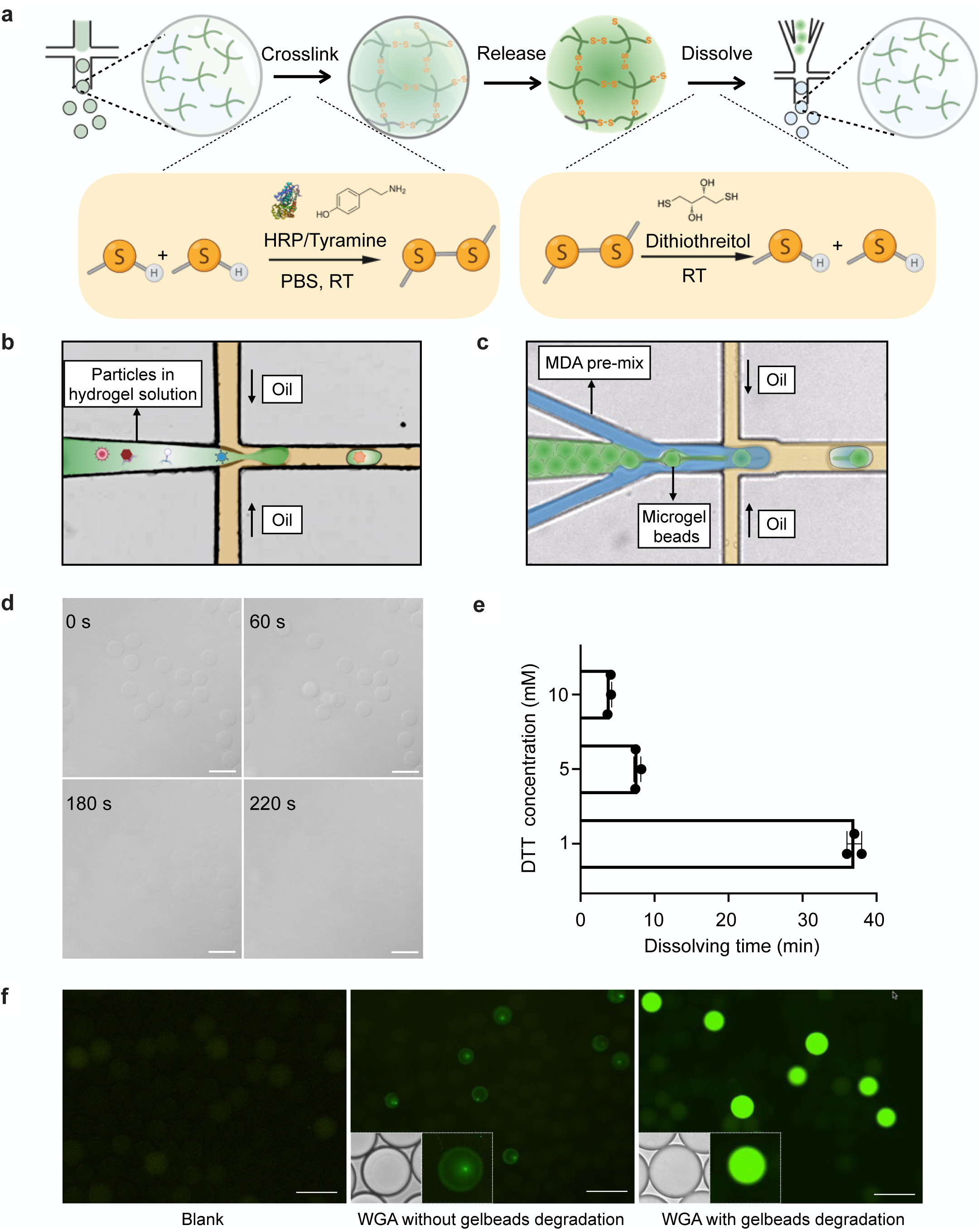
GSE-Seq, employed a strategy based on degradable hydrogel, is highly compatible to a variety of downstream analysis. **a,** The gelation and gel degradation in droplet. Orange box represents the chemical reactions of gelling and dissociation of hydrogel. **b-c,** Chip design for hydrogel solution encapsulation (**b**) and hydrogel beads degradation(**c**)**. d,** Degradation time of the hydrogel was assessed through a time course experiment. Scale bars, 50 µm. **e,** Degradation time of the hydrogel at different DTT concentrations. **f,** In-droplet whole genome amplification under different conditions. Scale bars, 100 µm.

The encapsulated entities, such as a single bacterium or virus particle, underwent lysis, and their DNA was purified within the gel beads using multiple steps under harsh conditions (Figure S2b). After preparing purified genomes, WGA was conducted by co-encapsulating the WGA reagents containing Dithiothreitol (DTT) and hydrogel beads in droplets (Figure 2a,c, Video S3). An addition of 10 mM DTT was shown sufficient to degrade the hydrogel in the water-in-oil droplets in 10 mins (Figure 2d, e). The WGA efficiency was significantly increased compared to the undissolved gel beads (Figure 2f). To reduce non-template amplification and improve uniformity in WGA^11^, we used the primase-based WGA instead of the conventional hexamer-based method. Easy deactivation of the primase reduces interference in the following library preparation. Developing a well-designed degradable hydrogel, combined with droplet microfluidics, has the potential to revolutionize droplet-based single-cell analysis by making it applicable to a wider variety of cell types. This is especially important for cells that are challenging to lyse, such as those found in microbiomes, plants, algae, and other organisms.

### Novel in-droplet library strategy for long-read sequencing

After WGA, library preparation, typically involving multiple enzymatic and purification reactions, can be a significant challenge if all processes are conducted entirely inside water-in-oil droplets. To enable library preparation, including the most challenging step of barcoding the single amplified genome (SAG), directly in water-in-oil droplets, we developed a novel enzyme mixture that achieved fragmentation and end-repair using the same enzyme mixture. To generate sufficient DNA fragments for barcode ligation, T7 Endonuclease I was employed to fragment highly branched WGA products, followed by adding a non-templated dA tail to the PCR products via Hotstart *Taq* DNA polymerase (Figure 3a). To initiate DNA fragmentation, the reaction was initially performed at 37°C to ensure optimal endonuclease activity. Meanwhile, the polymerase activity was effectively inhibited by the reversible binding of an aptamer at temperatures below 45°C. Subsequently, the endonuclease was deactivated by raising the temperature to 72°C, and the polymerase activity was activated. This temperature switch strategy enabled simultaneous DNA fragmentation and end-repair within the same droplets, resulting in DNA fragments with barcode attachable ends (Figure S3a).

**Figure 3.**
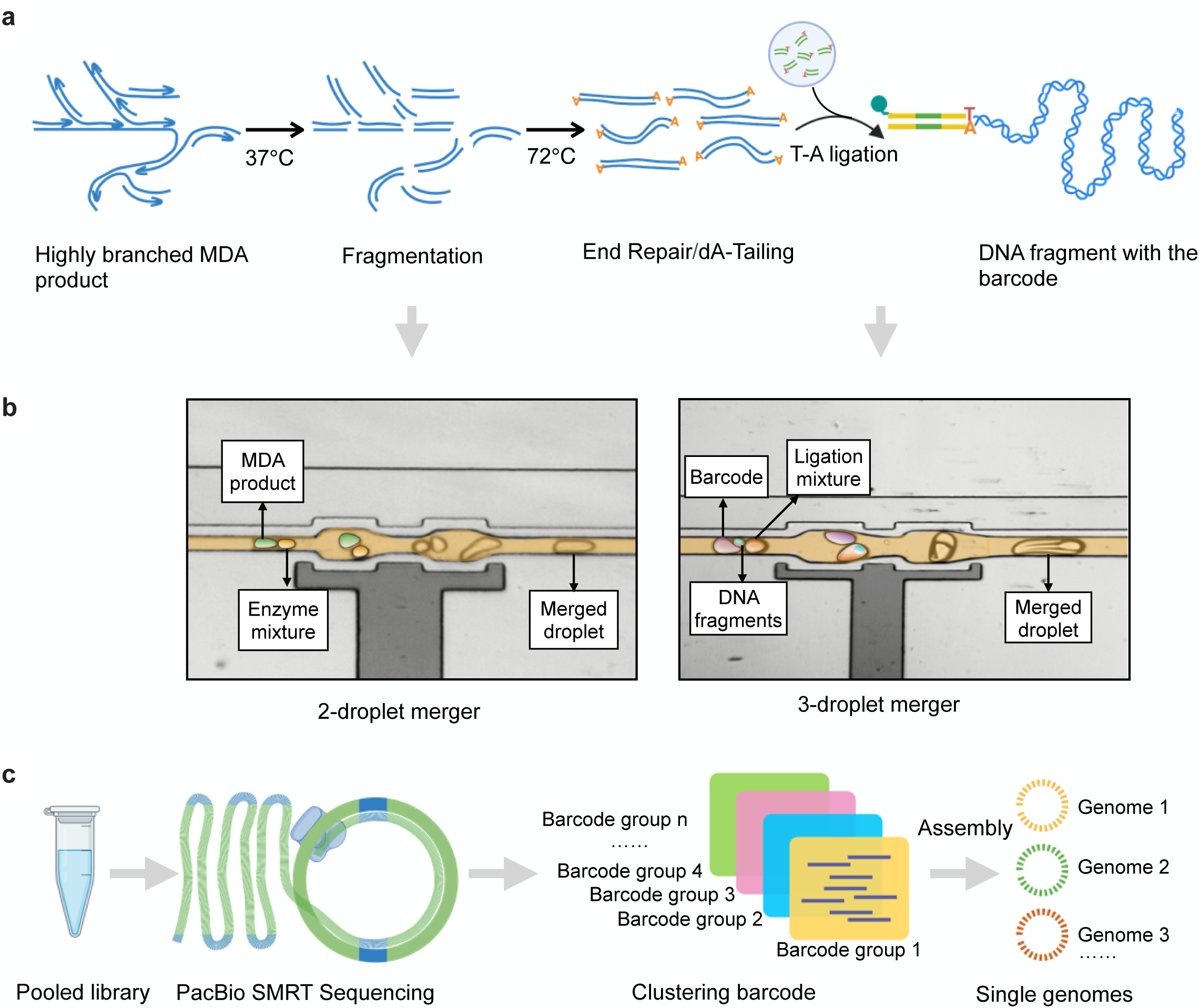
In-droplet library strategy for long-read sequencing. **a,** Single amplified genome barcoding through T-A ligation. **b,** Simultaneous fragmentation and end-repair via one-step 2-droplet merging (left) and T-A ligation via 3-droplet merging (right). **c,** Workflow to obtain the genome sequence of single entities from a pooled library.

The droplets containing highly branched WGA products and enzyme mixture were mixed by the specially designed 2-droplet merger (Figure 3b, Video S4). Following that, the droplets containing the attachable WGA products, barcodes, and ligation reagents were mixed in meticulous control 1:1:1 ratio using the 3-droplet-merger (Figure 3b, Video S5). Utilizing the optimal temperatures for the employed enzymes, in-droplet fragmentation and T-A ligation took place sequentially in a well-controlled manner. The resulting barcoded genomic DNA was then pooled by breaking the droplets. After purification and DNA repair, the barcoded DNA library was then amplified to generate sufficient material for sequencing (Figure S3b,c). To the best of our knowledge, this is the first strategy to achieve barcoded T-A ligation in droplets, a powerful approach for DNA library preparation in genetic utility. As most DNA manipulations, such as genome cloning and library preparation, rely on T-A ligation^12^, our new in-droplet TA ligation workflow can be applied in various applications.

Current single-cell analysis methods are often limited to short-read sequencing, which is more straightforward and established but presents enormous challenges in achieving accurate and complete genomes^13^. Our library has been designed to ensure compatibility with various commercially available long-read sequencing platforms, such as Nanopore and PacBio. To begin with, we conducted a pilot run using the Nanopore portable MiniON sequencer. The PacBio circular consensus sequencing (CCS) platform was chosen to generate high-fidelity (HiFi) reads to achieve higher read quality. Following sequencing, we can distinguish the raw reads from the pooled library into individual entities using their unique barcode identities (Figure 3c). In summary, the novel designs mentioned above offer a powerful and versatile solution to establish a comprehensive and generic workflow for single-entity sequencing. These advancements significantly enhance efficiency, simplify the process, and ensure compatibility across various applications, making generic single entity sequencing available.

### Characterization of GSE-Seq by a mock community

We applied GSE-Seq to analyze a mock viral population of known phages to assess our platform’s performance. The HiFi SAGs library generated by GSE-Seq produced high-quality reads (96.5 % > Q30) with an N50 1.7 kb, which is about six times of current short-read method (Figure S5a). External contamination poses a substantial challenge in WGA for single-entity sequencing. Traditional approaches, such as manual picking or fluorescence-activated cell sorting (FACS), have exhibited lower microbial positivity rates, with only 56.4% and 65.6%, respectively^14^. Of all the reads analyzed, 95.3% were aligned to the virus reference genomes. In contrast, less than 0.05% of reads were mapped to the human genome (Figure 4a), demonstrating the successful contamination prevention measures and utilization of droplets as isolated reaction chambers in mitigating environmental contamination. Unaligned reads may be due to chimeric reads generated during whole-genome amplification, library amplification, or sequencing.

**Figure 4.**
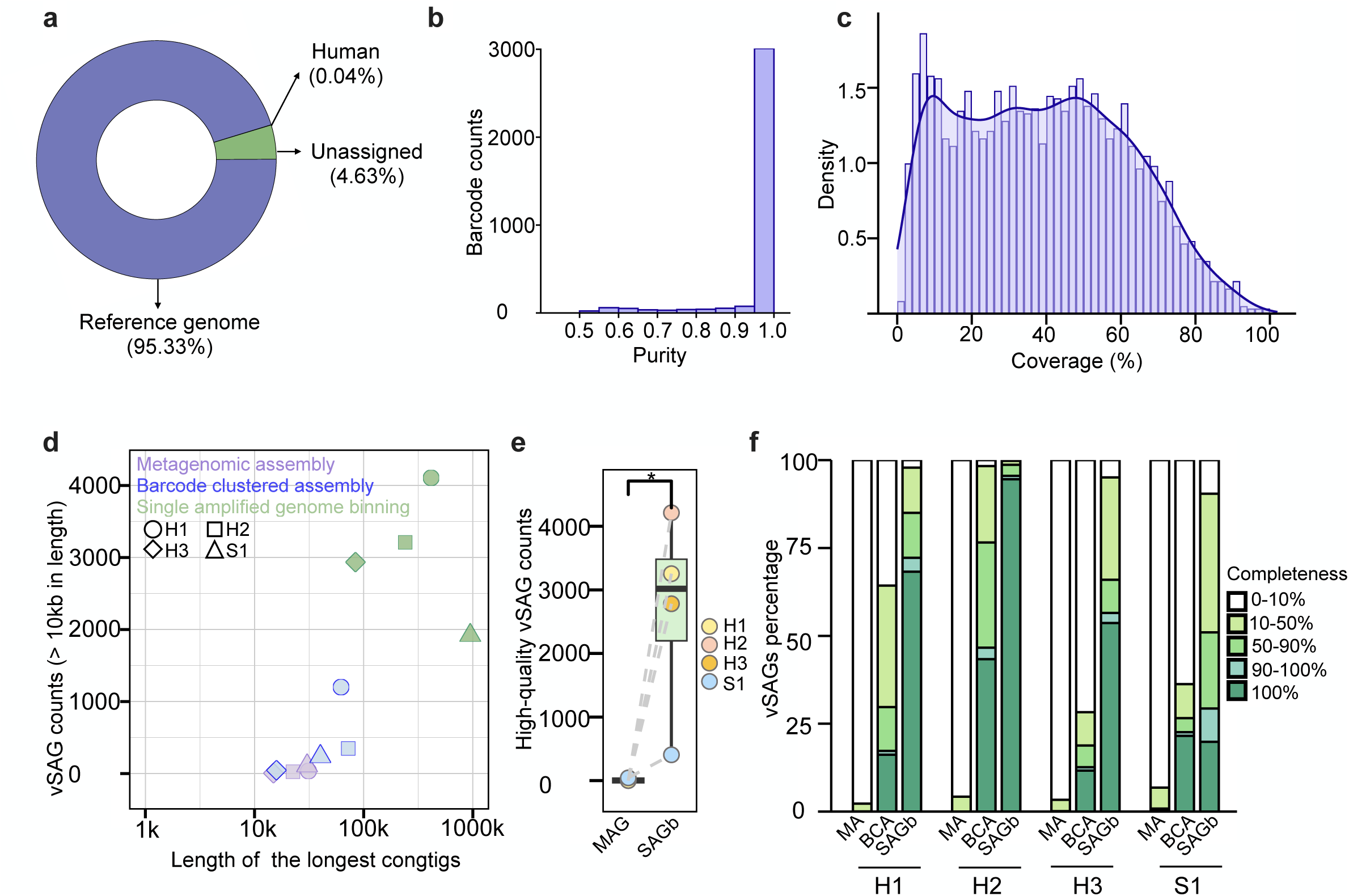
Validation the performance of GSE-seq. **a-c:** Mapping ratio (**a**), purity (**b**), and genomic coverage (**c**) of the viral mock community. **d,** Assembly statistics for metagenomic assembly, barcode clustered assembly, and single amplified genome binning. **e,** Number of high-quality genomes obtained using different assembly methods. *, p < 0.05. **f,** Completeness distribution of fecal and marine viral genome assembly using different methods. MA: metagenomic assembly; BCA: barcode clustered assembly; SAGb: single amplified genome binning.

Barcodes were used to group reads, resulting in 3,564 barcode clusters (Figure S5b). Of these clusters, 98.8% (3523/3564) showed less than 1% external contamination (Table S2). We also mapped all reads from each barcode cluster to their reference genomes to assess the cross-contamination between barcode clusters. The high-purity barcode clusters, which accounted for 85.2% (3,036/3,564), showed less than 1% external contamination and a purity greater than 95% (Figure 4b and Table S2). The coverage of single cells has posed a long-standing challenge. For instance, the Tn5-based single-cell amplification strategy often yields a maximum coverage of less than 50%, rendering a 50% dropout of the fragments^15^. By implementing the GSE-Seq featured in improved WGA and library preparation strategy, 23 viral Single amplified genomes (vSAGs) were observed with more than 90% genome coverage, while 998 vSAGs (32.8%) exhibited a coverage ranging from 50% to 90% (Figure 4c, Table S2). However, the coverage of vSAGs also revealed bias, which may result from stochasticity in lysis, genome amplification, and library amplification bias (Figure 4c, S4b).

### Improved genome assembly by GSE-Seq

Viruses, encompassing various genomes, were used as a model to showcase the generic applicability of GSE-Seq across different genome types. On the other hand, bacteria were employed as a prime example of hard-to-lyse organisms. Gut and environmental samples, containing a high diversity of biological entities, were evaluated to demonstrate the versatility of GSE-Seq for real-world applications. In short, GSE-Seq was employed to analyze viral and bacterial communities in the human gut and marine samples, exploring single virus sequencing (SV-seq) and single bacteria sequencing (SB-seq).

Genome assembly in complex communities poses a persistent challenge. Applying single-cell assembly helps resolve mis-assembly and fragmented assembly commonly encountered in metagenomic assembly, mainly due to shared and repetitive regions^16^. We observed a significant improvement in the assembly after barcode read clustering (Figure 4d). For virus samples, the length of the longest contigs improved 1.9-fold (Figure S6a). The count of contigs greater than 2,000 bp improved 26-fold (Figure S6b). Additionally, we observed a 15.1-fold improvement for read counts larger than 10,000 bp (Figure S6c).

Binning often involves classifying DNA sequences into clusters that potentially represent an individual genome or genomes from closely related organisms^17^. Single-genome sequencing offers an additional advantage in the form of barcode clusters, which facilitate the "binning" of contigs into longer scaffolds^18^. Notably, the longest scaffolds in the gut virus sample was 5.1 times larger than the contigs after barcode binning (Table S3). Interestingly, the marine sample exhibited a more significant increase, with a 23.6-fold increase of about four times higher than the gut sample (Table S3). Furthermore, binning short contigs into longer scaffolds decreased the number of contigs longer than 2,000bp (Figure S6b), which showed that short contigs had been grouped into fewer longer scaffolds. The number of scaffolds over 10,000 bp increased by 21.2-fold (Figure S6c, Table S3). We proceeded to evaluate the assembly performance of this methodology for larger genomes – specifically gut bacteria. The results mirrored the previous findings, highlighting that bacterial single amplified genome binning (SAGb) dramatically enhanced assembly length (Table S3).

As anticipated, SAGb exhibited considerable potential in the recovery of high-quality viral genomes in comparison to metagenome assembly. The binning procedure notably yielded more complete (completeness = 100) viral genomes (Figure 4e, Figure S6d). With 100% completeness in gut viral Single amplified genomes (vSAGs), SAG binning delivered a staggering 1736.5-fold increase, while marine samples saw a 9.5-fold surge (Table S4). Fragmented contigs were combined to form a more complete and high-quality (completeness ≥ 90) genome via barcoded clustered assembly and single amplified genome binning (Figure 4f, S6d). In this research, we showcased that the genome assembly could be significantly enhanced merely by partitioning complex metagenome reads without binning the single virus read. Moreover, the assembly process could be further amplified by binning individual viruses or bacteria scaffolds within a single genome.

### GSE-Seq uncovered the dark matter in the gut and marine viral genomes

The current SV-seq relies exclusively on flow cytometry sorting for VLP isolation^18^, which results in labor-intensive operation and prohibitive cost. Consequently, in previous single virus genomics investigations, each study only obtained dozens of viral genomes, representing merely a minute proportion of the vast viral diversity^4,19^. We recovered a total of 15,710 vSAGs from 3 healthy donors (Table S5) and 2,467 vSAGs from marine sediment (Table S6). Among these, 59.6% (10,828) demonstrated 100% completeness, and 63.0% (11,443) showed larger than 90% completeness (Figure 5a). We successfully acquired vSAGs encompassing all types of DNA, including dsDNA,ssDNA, and both linear and circular forms. Approximately 52.2% of the vSAGs from the human gut (H1: 44.4%, H2: 90.1%, H3:29.0%) and 12.8% from marine sediment were identified as ssDNA (Figure 5b). In addition, our analysis uncovered the presence of complete circular viral genomes among the vSAGs, with 5.4% observed in the human fecal sample and 2.2% in the marine sediment (Figure 5c). This highlighted our method’s ability to obtain the entire circular genome, although the proportion might be underestimated due to incomplete assembly.

**Figure 5.**
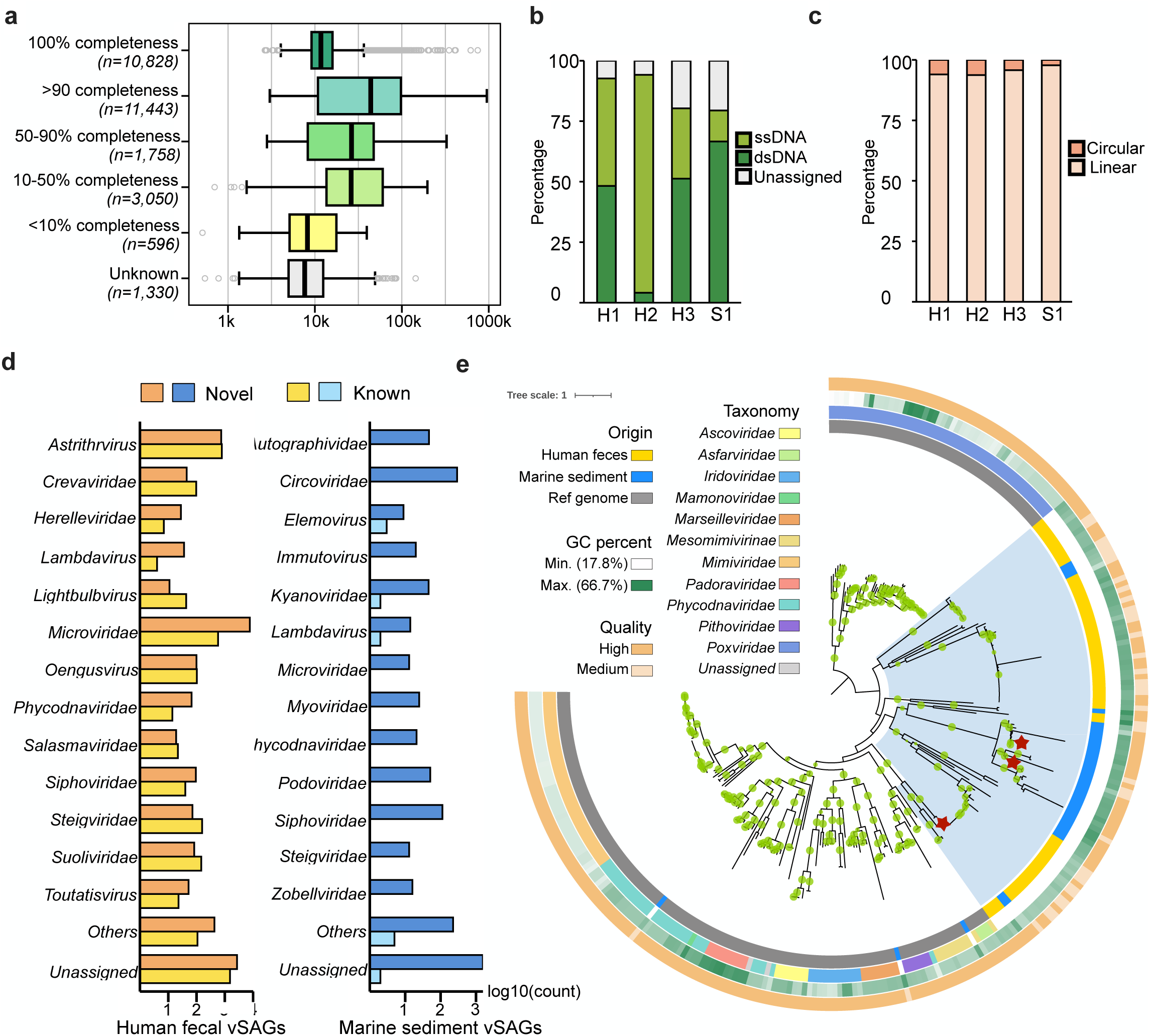
Application of GSE-seq to viral communities from human feces and marine sediment. **a**, Distribution of estimated genome completeness and length of viral single amplified genomes (vSAGs) into quality tiers. **b**, Baltimore classification of vSAGs from human feces and marine sediment. **c**, Complete circular genomes detected by GSE-seq. **d**, Distribution of novel vSAGs across different families or floating genera. **e**, Maximum-likelihood tree of 100 putative NCLDV vSAGs together with isolate NCLDV reference genomes. Tree annotations from inside to the outside: (1) origin, (2) taxonomy, (3) GC content and (4) genome quality. Green dots on the branches represent bootstrap support ≥ 70%. Blue shadow shows the main clades distribution of the putative NCLDV SAGs in this study. Red stars denote 3 vSAGs with genomic structures shown in Figure S7.

Viral metagenomics has proven to be a powerful tool for discovering new viruses. However, approximately 60-95% of viruses remain uncharacterized, earning them the "dark matter"^20^. Thus various viral catalogues were compiled from numerous metagenomes to enlarge reference databases^21,22^. In this study, we compared our vSAGs to publicly available databases and found only 23.1% of gut vSAGs (H1: 24.5%, H2: 10.9%, H3: 31.5%) exhibited significant similarity at the species level (Figure 5d, Table S5). Notably, the identification rate was even lower for marine vSAGs, with only 9 (0.4%) matches identified when compared to the NCBI RefSeq and the recent GOV 2.0 metagenomics database (Figure 5d, Table S6), indicating the sea holds untapped viral diversity with novelty, particularly the East Asian sea. These findings highlighted the powerful application of single virus sequencing in uncovering viral "dark matter" in humans and the sea.

Only 73.1% of human fecal vSAGs (Table S5) and 36.5% of marine vSAGs (Table S6) could be annotated at the family or floating-genus level. *Microviridae*, one of the most stable colonizers of the human gut^23^, was predicted to account for the vast majority of ssDNA viruses in these three individuals (Figure 5d, Table S5). In contrast, ssDNA viruses in the sea sample were mainly composed of *Circoviridae* (Figure 5d, Table S6), which exists in numerous animal hosts and has recently been found in oysters of *C. hongkongensis^24^*. Meanwhile, most of the dsDNA virus belonged to *Caudoviricetes* class, mainly families *Siphoviridae*, *Podovirida, Myoviridae, and Steigviridae* (Figure 5d, Table S5, S6). The smallest genome we have found with 100% completeness was 2655bp, which belonged to the *Circoviridae* family (Table S6).

Nucleocytoplasmic large DNA virus (NCLDV) is a recently discovered group of large, complex viruses with genomes and structural features previously thought to be exclusive to cellular organisms, challenging our understanding of viral evolution and blurring the boundaries between viruses and cells^25^. Utilizing GSE-Seq, we could detect Nucleocytoplasmic Large DNA Viruses (NCLDVs). This enabled us to delve into their diversity, an aspect frequently overlooked by conventional metagenomic strategy due to these viruses’ larger genome size and scarcity^25^. We successfully identified 100 vSAGs possessing at least two Nucelocytoviricota marker genes (Table S7). The largest genome identified was 948 kilobases, marking the longest vSAG in our study. However, using a 0.22 μm filter during the VLP enrichment process might have excluded larger VLPs, potentially causing an underestimation of NCLDVs in the analyzed samples. The workflow can be easily modified for NCLDV targeting via VLP size selection. We construct an IQ-tree using the 100 putative NCLDV SAGs and reference genomes of NCLDV isolates (Figure 5e). Excluding three vSAGs which were placed within *Mesomimivirinae, Marseillviridae,* and *Phycodnaviridae*, the majority of putative NCLDV SAGs were clustered into independent clades with no isolate representatives (Figure 5e).

Meanwhile, our inferred NCLDV vSAGs showed a considerable divergence from the existing NCLDV viruses catalogued in the Giant Virus Database (GVDB). Only a single vSAG exhibited more than 70% Average Amino Acid Identity (AAI) and 20% coverage with the reference genomes (Table S7). COG annotations disclosed that genes associated with DNA replication, translation, transcription, the metabolism of carbohydrates and amino acids, and energy production and conversion were abundantly present in the putative NCLDV SAGs (Figure S7, Table S8).

Despite being highly abundant, the crAssphages with their significant heterogeneity and larger genome size -- remained a mystery until 2014, when cross-assembly methods were used to describe them^26^. However, GSE-Seq would be an invaluable tool for retrieving crAssphage from complex communities. In the human samples we analyzed, more than two conserved genes^27^ of crAssphage were detected in 92 vSAGs from H1 and 88 vSAGs from H3 (Table S9). Interestingly, we also detected 19 crAssphage signals from a marine sediment sample, which is, to our best knowledge, the first reported marine sample in Hong Kong. Given that crAssphage is believed to be an excellent marker for human fecal contamination^28^, which suggested the potential human-fecal contamination of the Hong Hong Coast ecosystem may have occurred, as the marine sediment sample collected near the Hong Kong Airport (Figure S8). In summary, GSE-Seq allows us to detect all types of DNA, spanning from the minimum of 2.6Kb to the largest 948Kb NCLDV and even the hyper-variable crAssphage.

### GSE-Seq further uncovered the diversity of gut bacteria

We examined a human gut sample to assess the effectiveness of GSE-Seq in addressing large and difficult-to-lyse bacterial genomes, and we managed to recover 23,266 bacterial single amplified genomes (bSAGs) (Table S10). Impressively, we were able to attain 2 bSAGs with a completeness of 90%. Additionally, we successfully produced 81 medium-quality bSAGs, showcasing completeness levels between 50% and 90%, each with a quality score of 10 or above (Figure 6a, Table S10). These 83 bSAGs were identified as belonging to the phyla *Actinobacteria*, *Firmicutes*, and *Bacteroidetes* (Figure S9, Table S11), the dominant phyla in the human gut^29^. We also achieved 1210 low-quality draft genomes exhibiting completeness between 10% and 50% (Figure 6a). More than 70% of human gut microbial species are still uncultured^30^. Among 83 medium or high-quality bSAGs, 32 belonged to putative known species, such as *Bifidobacterium longum*, *Dorea formicigenerans*, *Gemmiger formicilis*, and *Prevotella copri* (Table S11). However, 51 bSAGs (61.4%) represented putative novel bSAGs (Figure S9, Table S11).

**Figure 6:**
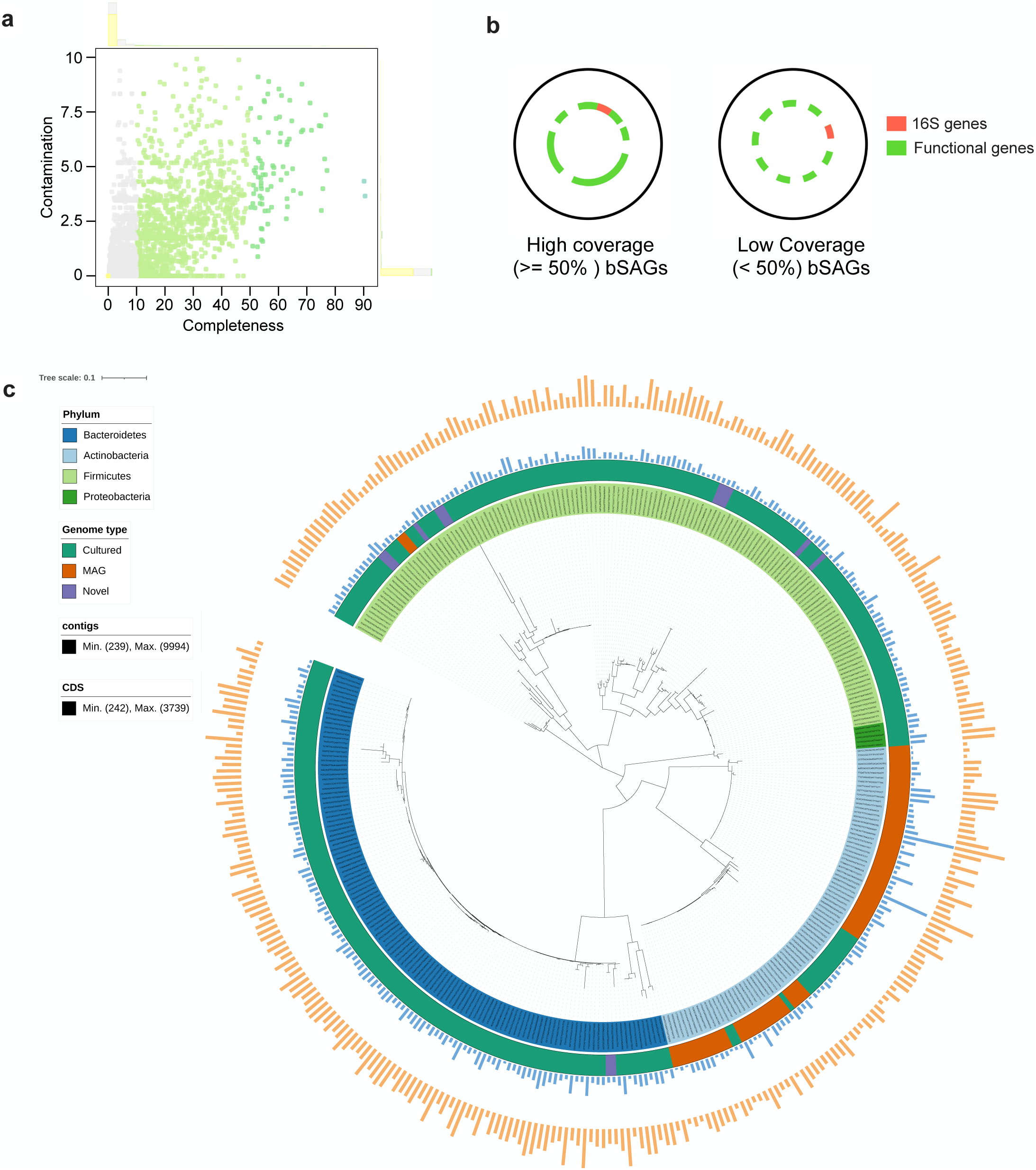
Validating GSE-seq in single bacteria genomes of human gut. **a**, Completeness of bacterial single amplified genome assembly (bSAGs). **b**, Analysis of 16S rRNA gene sequences and phylogenetic relationships. **c,** Taxonomic annotation was performed on the identified bSAGs. As a result, 360 bSAGs have near-complete 16S rRNA sequences (>1000bp). These bSAGs were taxonomically classified into 4 phyla, 6 classes, 6 orders, 9 families, and 27 genera. The tree annotations from inside to the outside are: (1) bSAGs colored by phylum annotation, (2) Genome type annotated by the UHGG database, including cultured strains, metagenome-assembled genomes (MAG), and putative novel genomes, (3) contigs lengths, and (4) CDS counts.

Due to the larger genomes of bacteria, low coverage remains a persistent issue in SB-seq. Two main methods, 16S ribosomal DNA (16S) and metagenomic-based phylogenetics, are conventionally used to investigate microbial diversity. Despite their limited coverage, bSAGs can serve as a feasible bridge between 16S sequencing and metagenomic-based approaches (Figure 6b). Primarily, this approach facilitates 16S phylogenetic analysis. In addition, it provides valuable insights into the function of genes within a specific taxonomic group - a challenge often met in metagenomic workflows due to the impediment of metagenome assembly by highly similar 16S sequences^31^. Therefore, despite the relatively low coverage for some bSAGs, it remains an invaluable tool for exploring the "microbial dark matter" and uncovering new insights into the vast diversity of uncultured microorganisms.

Among the all bSAGs, we found 2,946 that contained 16S rRNA gene sequences exceeding 500bp. After conducting a series of filtration and annotation processes, we discovered that 360 bSAGs had near-complete 16S sequences of over 1,000bp. These sequences spanned 4 phyla, 6 classes, 6 orders, 9 families, and 27 genera (Figure 6c, Table S12). Our 16S-based phylogenetic analysis revealed that 22.2% of the analyzed samples had less than 97% similarity to the 16S sequences of cultured species in the Unified Human Gastrointestinal Genome database. This suggested that these samples could potentially represent novel species identified via bSAGs. Interestingly, among these samples, 18.8% had only metagenome-assembled genomes (MAGs) as representative genomes (Figure 6c). We also discovered identified bacterial genera— including *Bifidobacterium, Collinsella, Clostridium XIVa, Dysosmobacter, Faecalibacterium, Lachnospira, Oscillibacter, Ruminococcus, and Ruminococcus2*—belonged to the Gram-positive which were typically more resistant to lysis (Table S12). This observation underscores the potential of GSE-Seq as an effective method for studying hard-to-lyse bacterial cells. Despite the inability to employ mechanical lysis methods such as bead beating in this study, the protocol could be amplified in efficacy by incorporating strategies like repeated in-situ freeze-thaw cycles or other alternative techniques^32^.

Prophages are integrated into the genomes of bacterial or archaeal hosts, which can make their identification more complicated. Short-read sequencing data may not provide the complete context of prophage-host genomes, leading to difficulties distinguishing between phage sequences and host genomic regions^33^. GSE-Seq, employing long-read sequencing at the single-bacterium resolution, emerged as a potent tool for prophage prediction. In this study, we detected 4,695 prophage signals across all bSAGs. Of these, 2,975 bSAG harbored at least one prophage, with a single bSAG containing up to 15 prophages (Figure S10a). Upon mapping the predicted prophages and bSAGs to their respective taxonomies, 1,494 high-quality pairs of prophages and bacteria were identified (Table S13). Remarkably, 97% of these prophages were classified under *Caudoviricetes*, the predominant class known for infecting bacteria. Furthermore, our analysis revealed a notable phage-host interaction rate of 28.1% for *Prevotella sp015074785* (Figure S10b). To probe the genomic similarity among prophages, we employed vConTACT to construct a gene-sharing network where viral clusters (VCs) approximate genus- or subfamily-level taxonomies based on sequences from the prokaryotic viral genomes in the RefSeq database. This analysis clustered the prophages into 193 VCs, of which only one included reference viral genomes, and 192 represented putative new viral genera (Figure S11). Notably, up to 45 prophage VCs were traced back to *Prevotella sp015074785*, followed by 27 prophage VCs originating from *Bifidobacterium longum* (Figure S11). With its capacity to capture host-phage interactions at the single-cell level, GSE-Seq is invaluable for unravelling the intricacies of viral infections, host immune responses, and their coevolutionary dynamics.

## Discussion

The Earth’s ecosystem harbors a vast spectrum of biological entities, the majority of which remain unexplored due to their immense diversity and heterogeneity^34,35^. To address this, we aim to create a high-throughput platform capable of capturing the remarkable biological diversity, particularly focusing on entities currently overlooked due to methodological constraints. To this end, we developed GSE-Seq, a novel platform designed to sequence the genomes of individual biological entities within complex communities on a grand scale.

In an effort to maximize versatility, we have reimagined our workflow and devised a new platform based on droplet microfluidics. This innovation incorporates a novel degradable hydrogel, an innovative barcoding strategy, a well-tailored library preparation strategy, and long-read sequencing. Integrating these elements within the GSE-Seq platform has facilitated the study of diverse biological entities, ranging from small viruses with genomes as concise as a few thousand base pairs to bacteria with mega-base-sized genomes that are notoriously difficult to lyse. However, with the integration of potent lysis methods, it also holds obvious potential for use in fungi and plant studies. It is feasible to increase the throughput by increasing the positive number of barcodes and MDA droplets via incorporating fluorescence-activated droplet sorting, thereby further reducing processing time, reagent consumption, and sequencing costs.

We have successfully applied this platform to viral and bacterial communities, yielding a plethora of individual genomes with high coverage. This output enables us to conduct in-depth downstream analyses, including discovering previously unknown genomes and identifying host-phage interactions. We have utilized GSE-Seq to discover new species in the gut and sea, and we foresee its broad applicability in various fields such as microbial ecology, environmental science, biotechnology, and human health. In conclusion, GSE-Seq represents a substantial step forward in high-throughput genomic sequencing at the level of single entities and holds immense promise for future biological and biomedical research.

## Author contributions

G.W., L.Z., Y.S., and F.Q. conceived the idea and methodology for the study and conducted the experiments and data collection. L.Z., G.W., Y.S., Y.D., W.L., C.L., and M.L. spearheaded the computational analysis. G.L., L.L., and X.B. contributed to the study’s methodology. G.W. and Y.S. led in manuscript drafting. L.Z., F.Q., Y.H., and J.Y. revised the manuscript. J.Y. and Y.H. supervised the entire study. All authors read and approved the final manuscript.

## Acknowledgements

We would like to express our gratitude to the Yu and Megan lab members for their helpful discussions. We also appreciate the valuable contributions of Chengchen Zhang, Xia Xu, Yong Feng, Fenfen Ji, Yufeng Lin, Danyu Chen, Guangyao Chen, Eagle Chu, and Zheng Zhang. This project was supported by National Key R&D Program of China (No. 2020YFA0509200/2020YFA0509203); Research Talent Hub-Innovation and Technology Fund Hong Kong (ITS/177/21FP); RGC Research Impact Fund Hong Kong (R4032-21F); Shenzhen-Hong Kong-Macao Science and Technology Program (Category C) Shenzhen (SGDX20210823103535016); Vice-Chancellor’s Discretionary Fund Chinese University of Hong Kong; Research Grants Council of the Hong Kong Special Administrative Region, China (project no.: CUHK 14207121 and 14219922), as well as the VC Discretionary Fund, provided by the Chinese University of Hong Kong (project #: 8601014).

---

**Figure S1.**
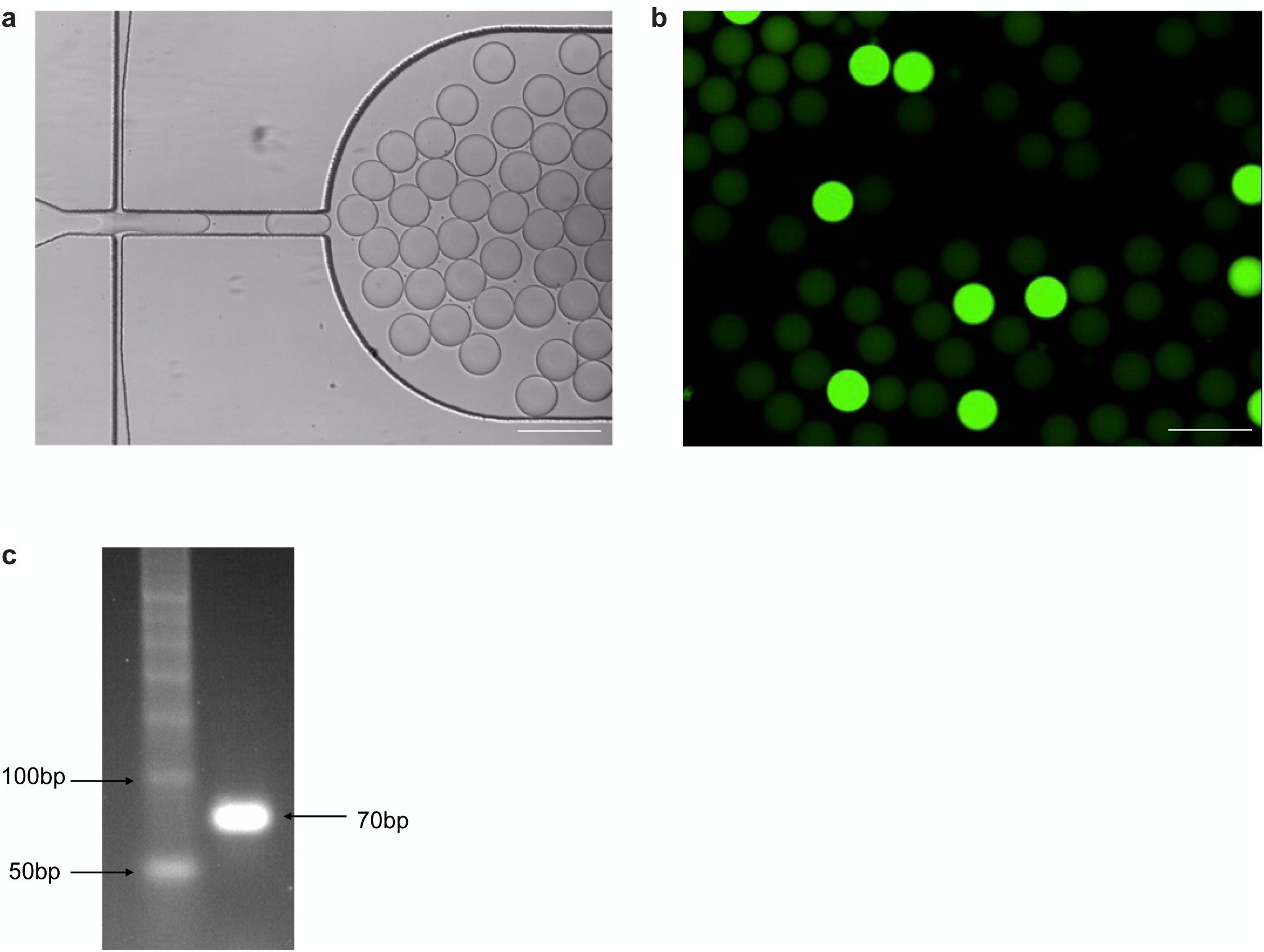
Monoclonal barcode amplification in droplets. **a**, Generating barcode on chip. Scale bar: 200 µm. **b,** SYBR staining of barcode droplets after PCR amplification. Scale bar: 200 µm**. c,** DNA electrophoresis of barcode PCR products after droplet breakage. The barcode fragment was ∼70bp.

**Figure S2.**
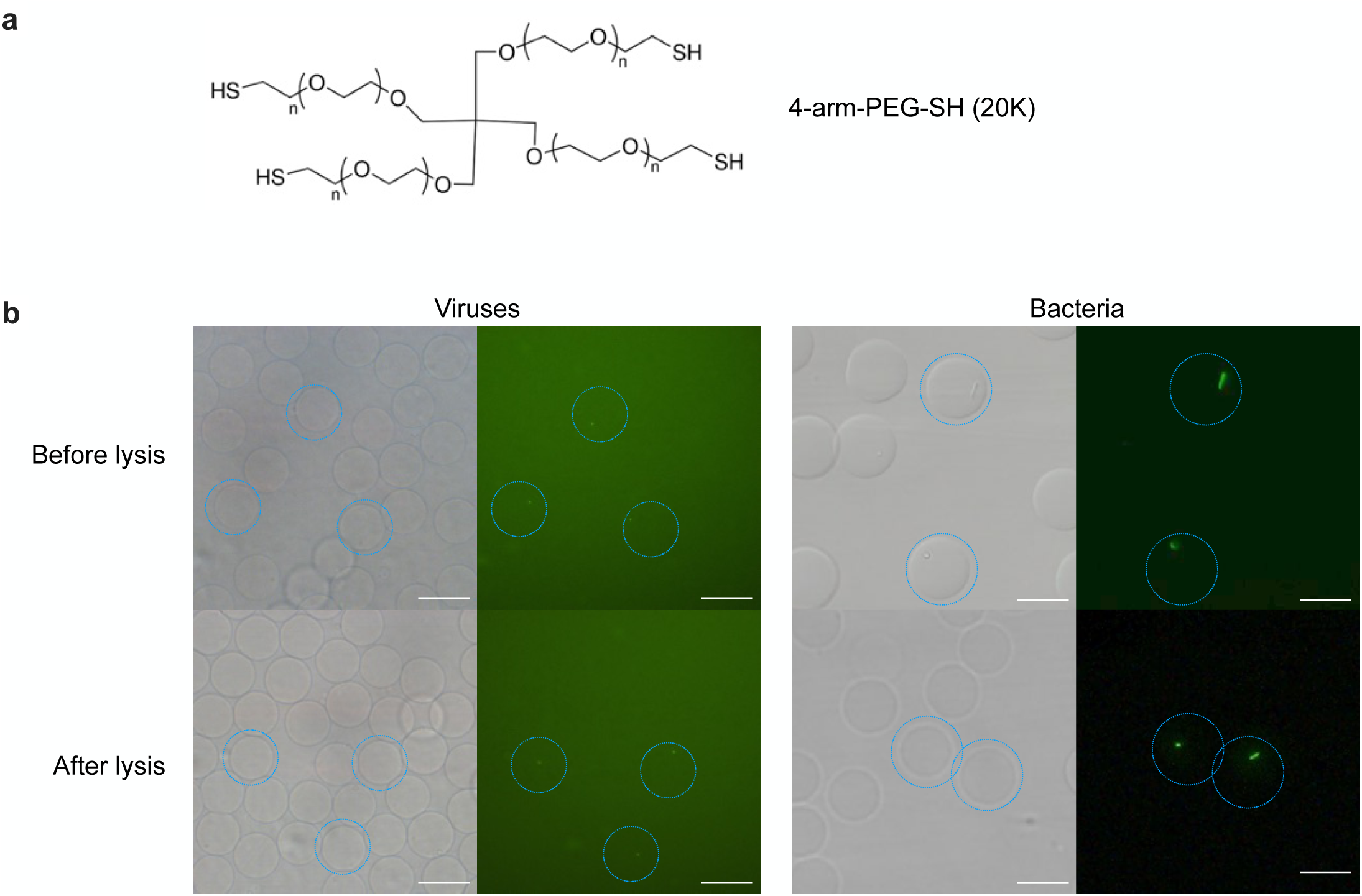
Degradable hydrogel for generic single entities lysis and purification. **a**, Chemistry structure of 4-arm-PEG-SH (20k). **b**, SYBR staining of viral (left) and bacterial (right) particles in hydrogel beads. Scale bar: 25 µm.

**Figure S3.**
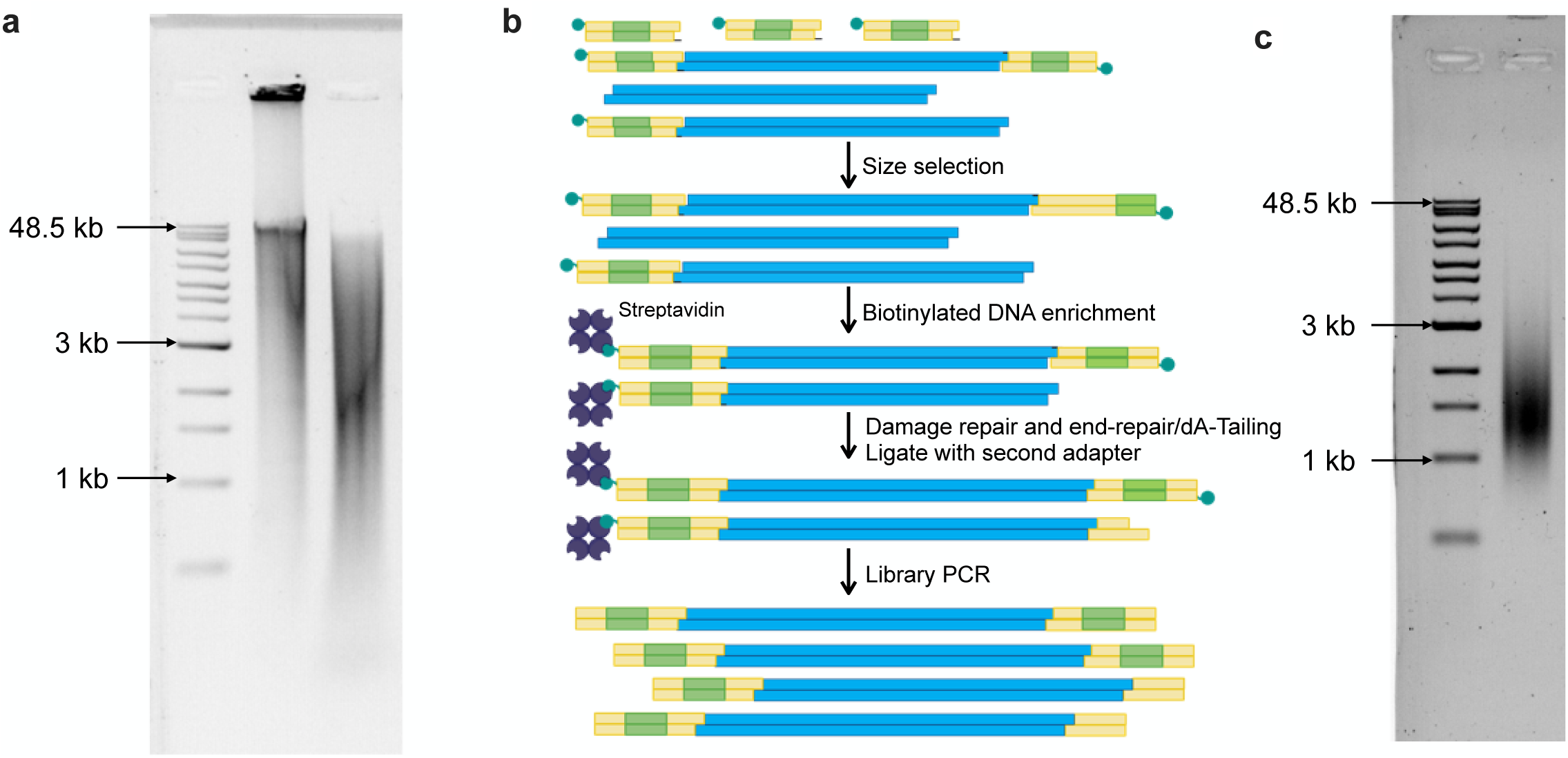
SAG library preparation. **a,** DNA electrophoresis before and after fragmentation/end-repair of highly branched whole genome amplification products. **b,** Schematic workflow of pooled library purification and amplification. **c,** DNA electrophoresis of GSE-seq library.

**Figure S4.**
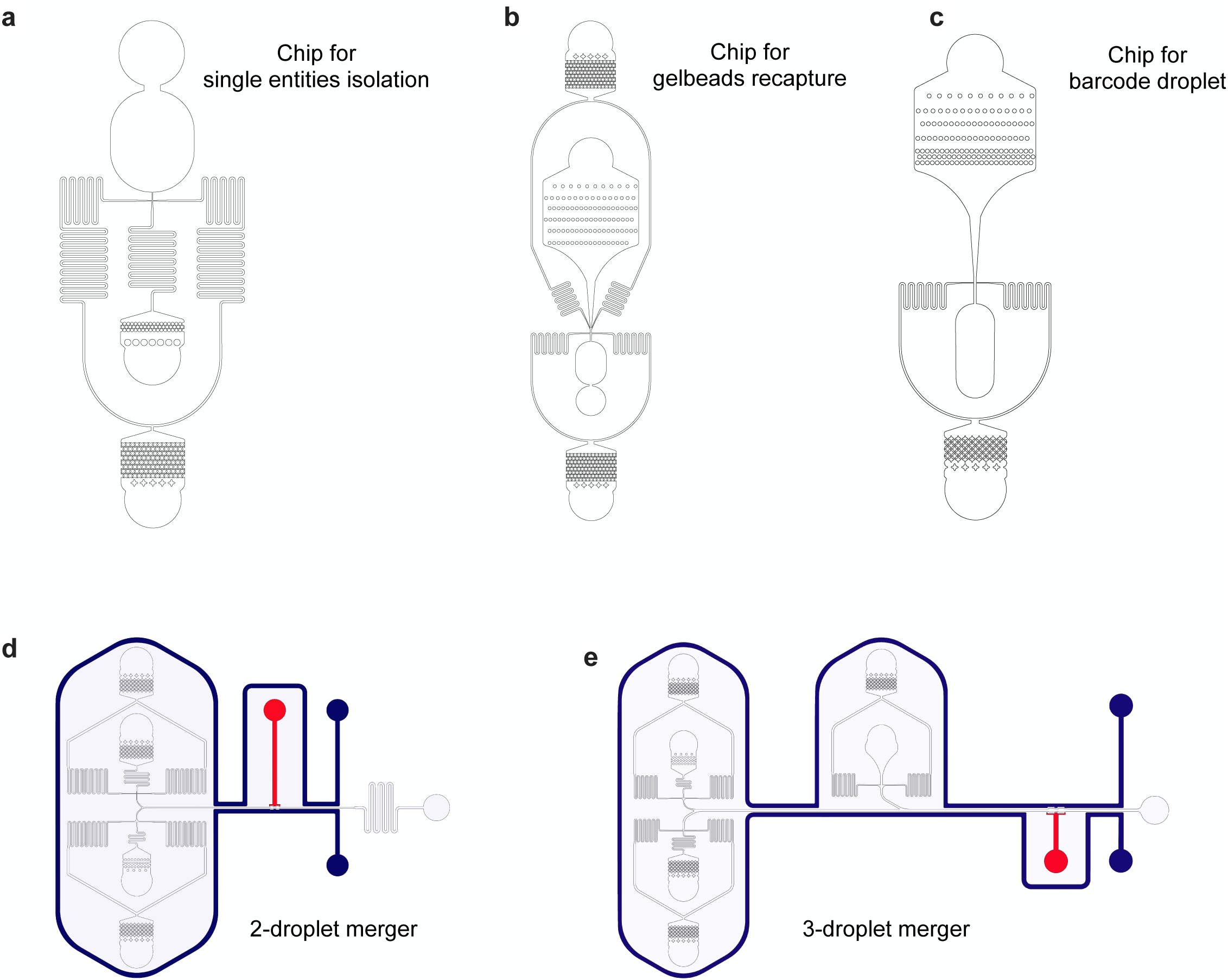
Microfluidic chips. **a**, Encapsulate single entities in hydrogel. **b**, Re-encapsulate microgels beads in WGA reagents. **c**, Generate barcode droplets. **d,** Merge 2 droplets to combine WGA droplets with T7 endonuclease I/ Taq polymerase enzyme mixture droplets. **e,** Merge fragmented WGA droplets with barcode droplets and ligation mixture droplets. The red color channel indicates the dead-end of the saltwater AC electrode, while the blue color channel indicates the ground electrode.

**Figure S5.**
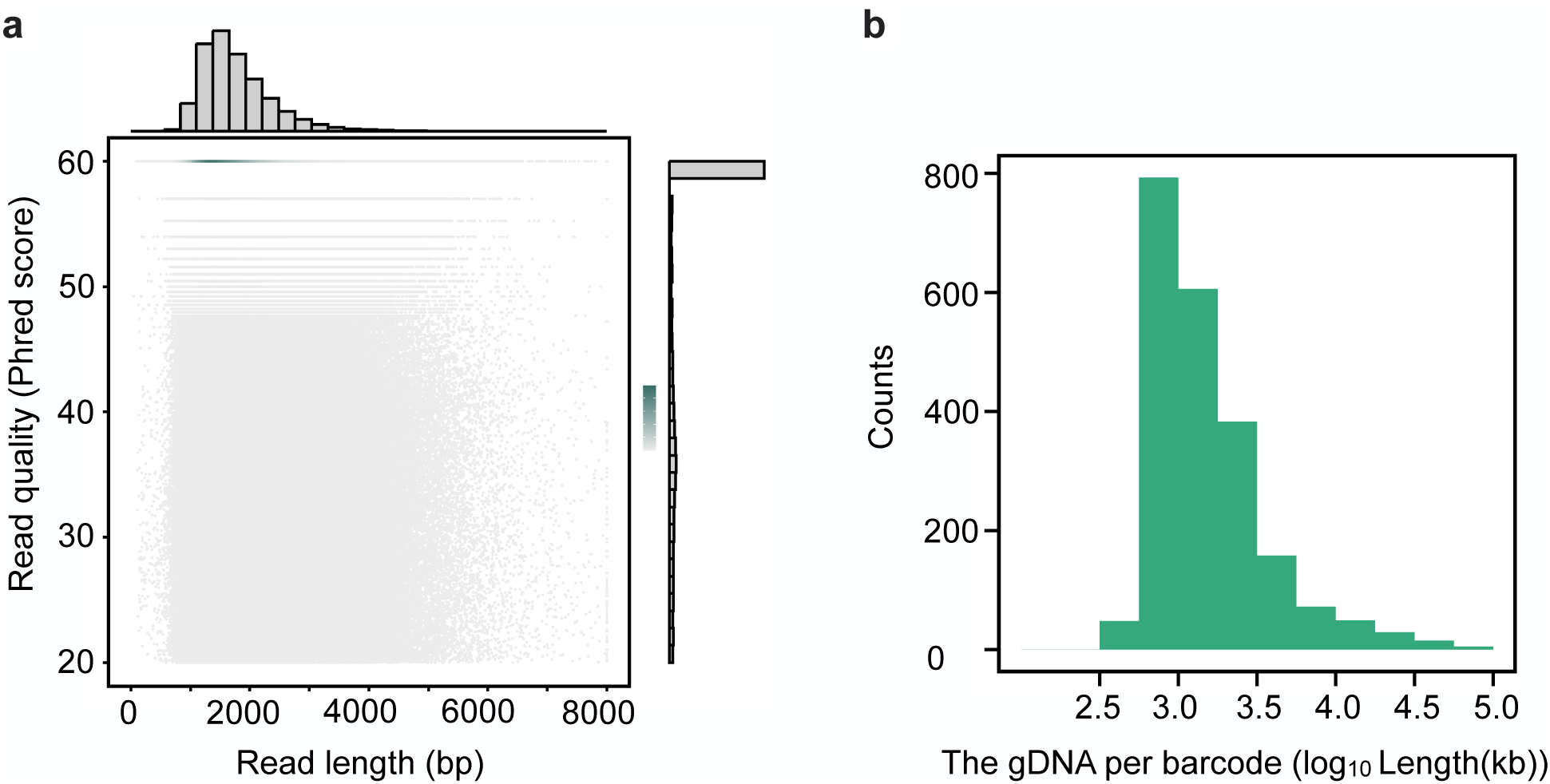
GSE-seq on viral mock community. **a,** Distribution of HiFi read quality and length. **b,** The gDNA distribution of each barcode cluster.

**Figure S6.**
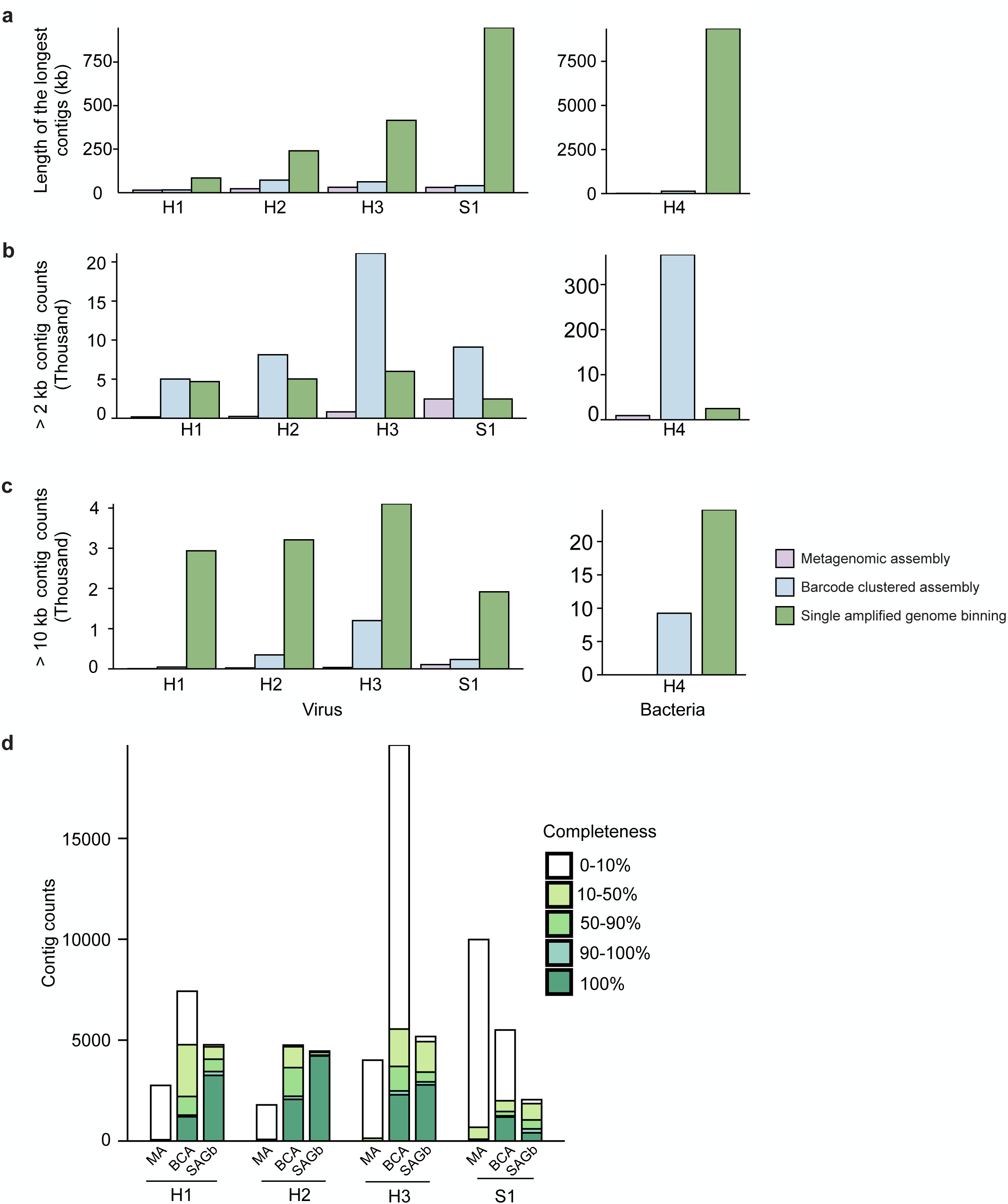
Assembly statistics of single amplified genomes from human feces and marine sediment. **a,** The longest contigs obtained through different assembly methods. **b-c,** The number of contigs longer than 2 kb (**b**) and 10kb (**c**). **d,** The number of contigs with different completeness. MA: metagenomic assembly; BCA: barcode clustered assembly; SAGb: single amplified genome binning.

**Figure S7.**
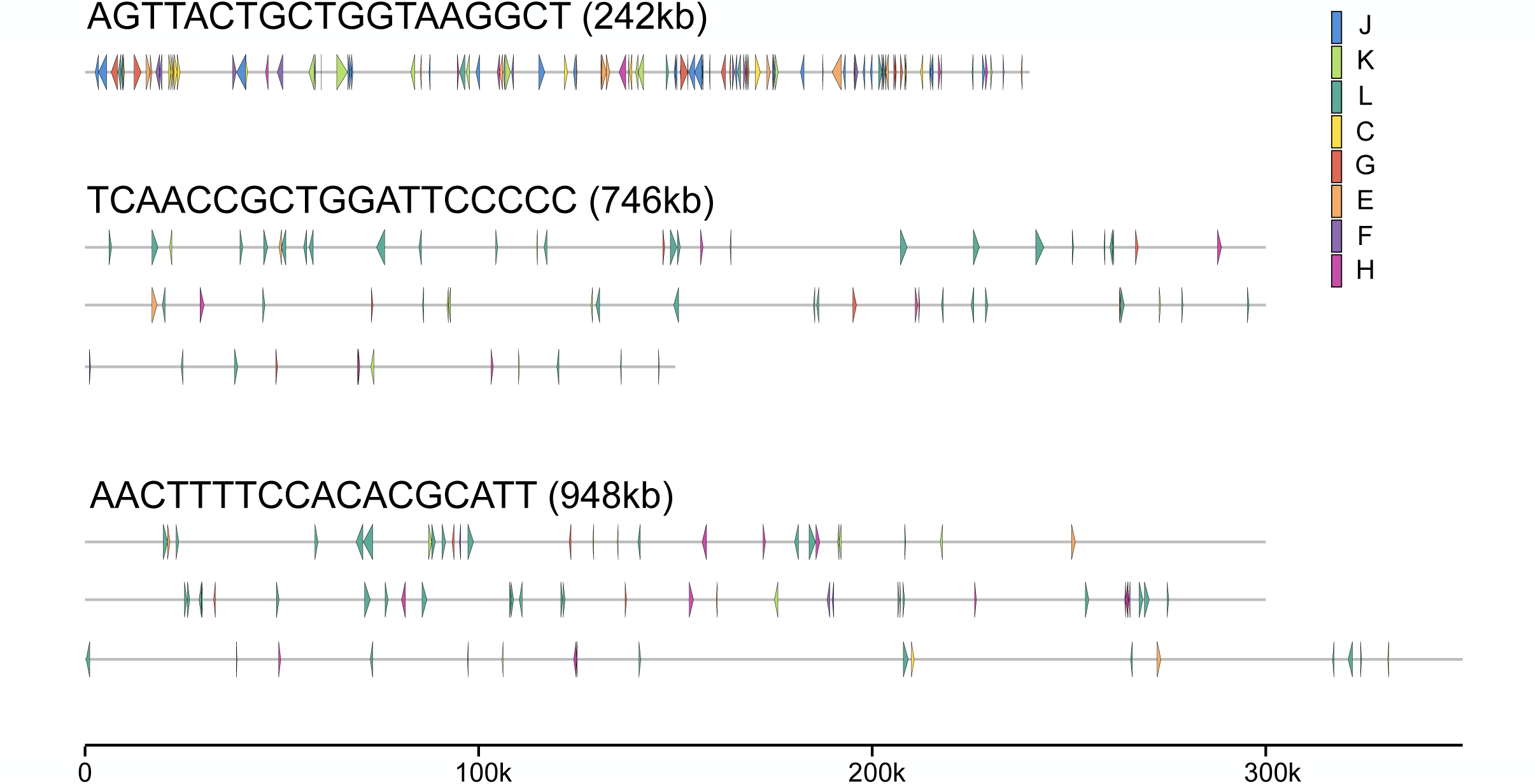
Presence of special functional genes in the putative NCLDV SAGs. Three representative NCLDV SAGs with over 100 known ORF gene functions are shown. Only genes involved in metabolism, DNA replication, translation, and transcription are presented. J: Translation, ribosomal structure, and biogenesis; K: Transcription; L: Replication, recombination, and repair; C: Energy production and conversion; G: Carbohydrate transport and metabolism; E: Carbohydrate transport and metabolism; F: Carbohydrate transport and metabolism; H: Coenzyme transport and metabolism.

**Figure S8.**
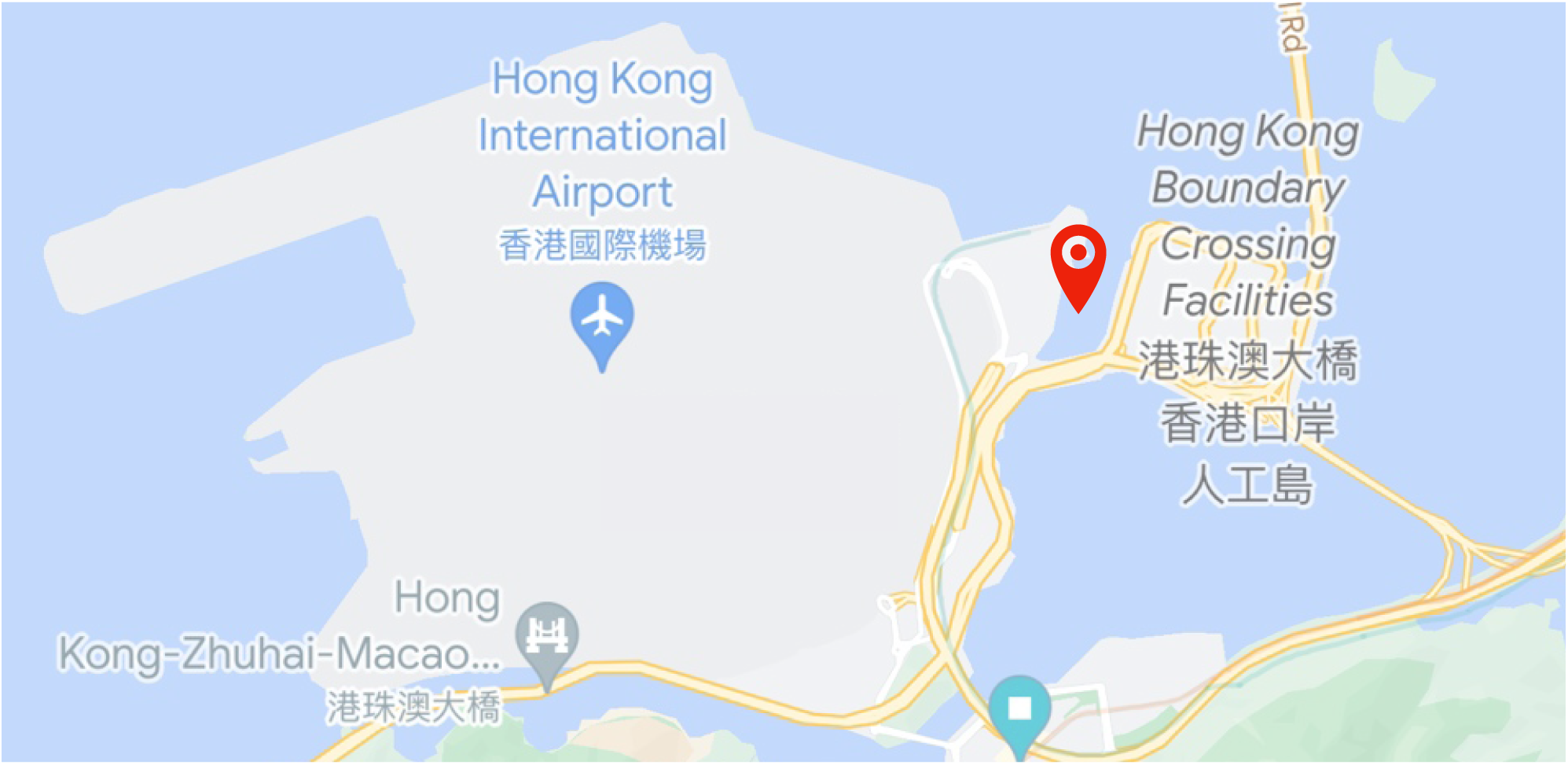
Extend information of viral sample for GSE-seq. The red label indicates the location for collecting marine sediment.

**Figure S9.**
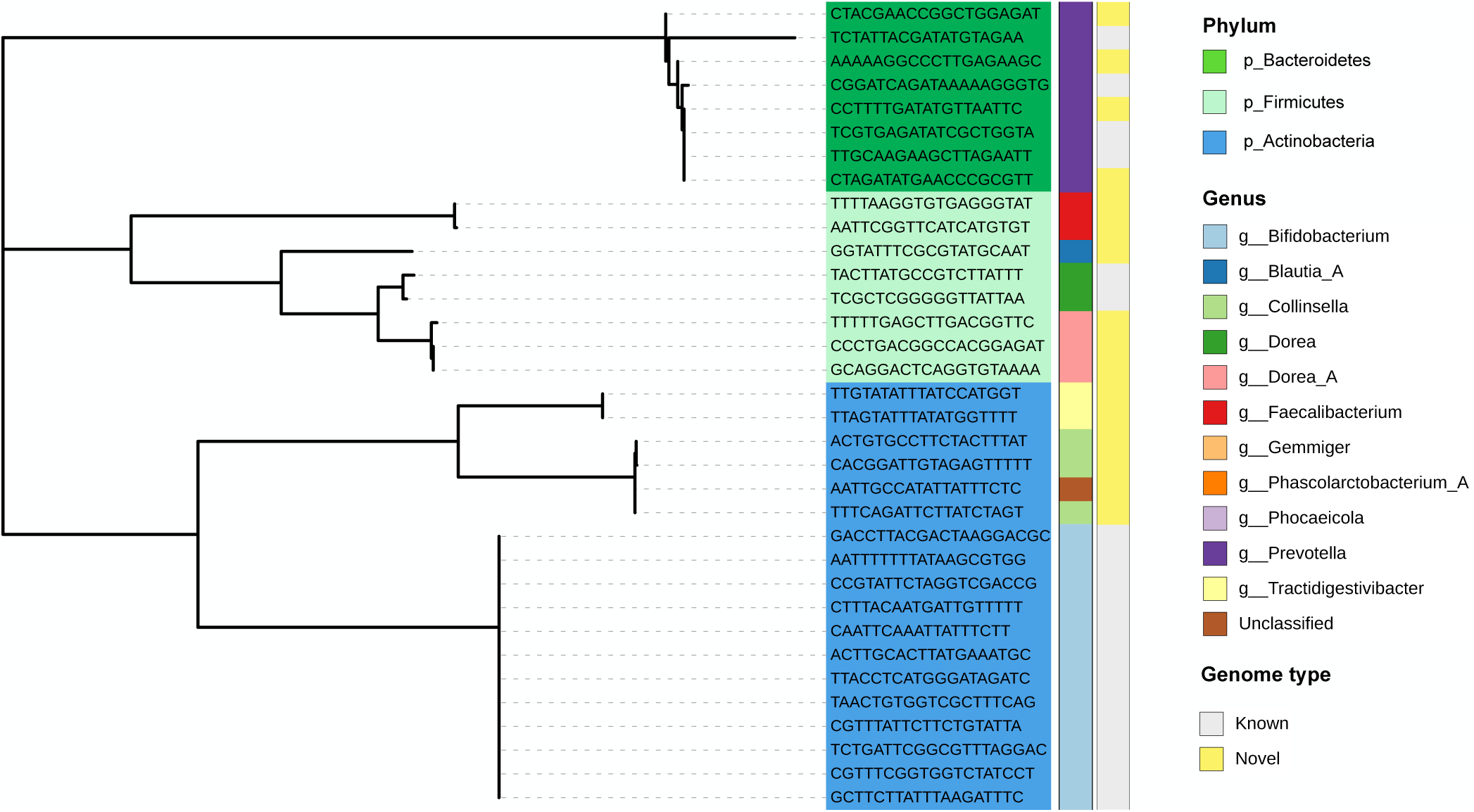
Phylogenetic tree of medium- and high-quality bSAGs. The label indicates the barcode of each bSAG, with text color-coded according to its phylum. The first column displays the genus annotation of each bSAG, and the second column presents the ANI-based genome information for each bSAG (known >= 95%, novel < 95%).

**Figure S10.**
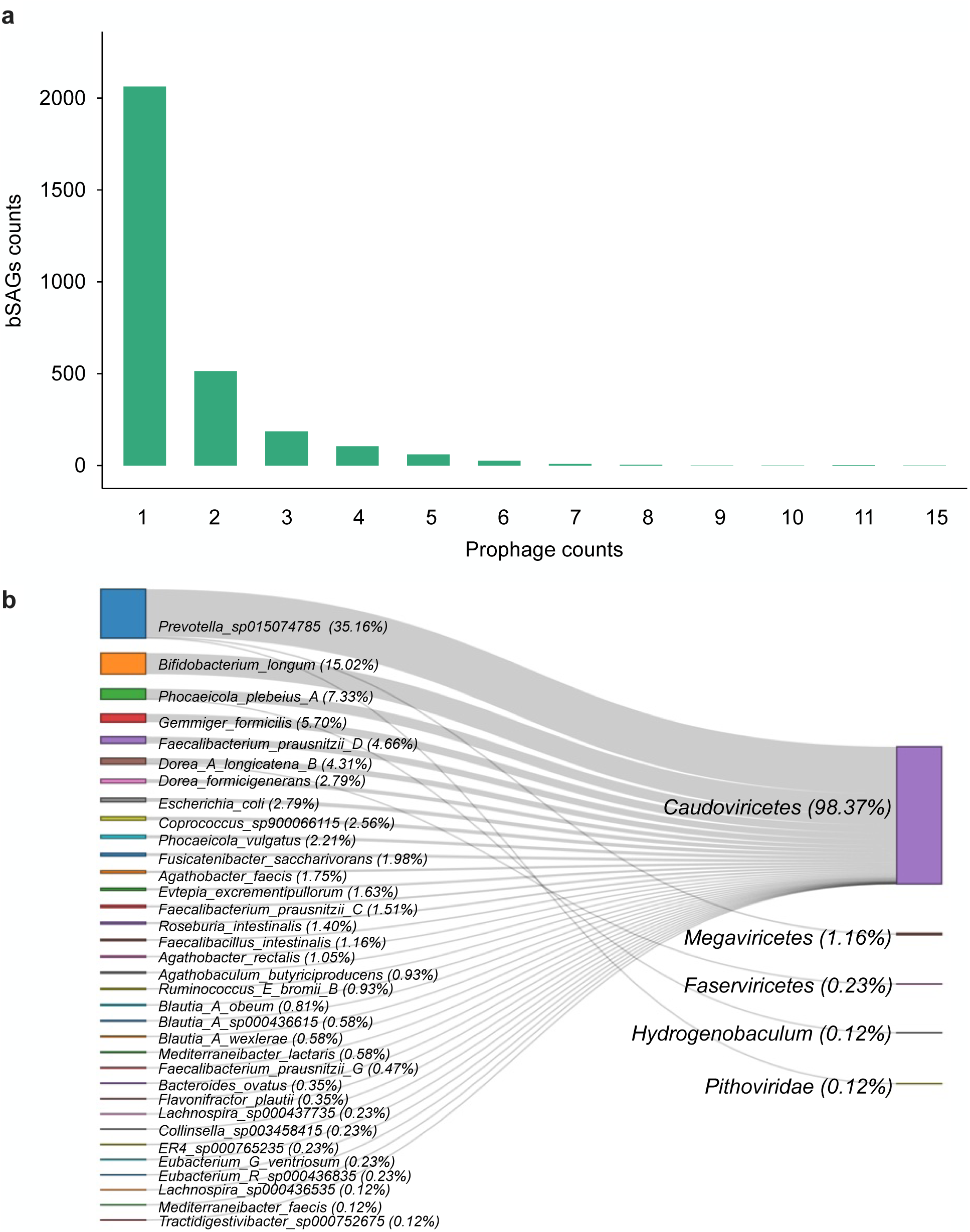
Prophage identified from bSAGs. **a**, Number of prophages identified from each bacterial SAG. **b**, Prophage-host pairs determined by analyzing prophage and host phylogeny. Prophages were grouped at the class level and bacteria at the species level, with each side showing the percentage of class/species among all the sequences

**Figure S11.**
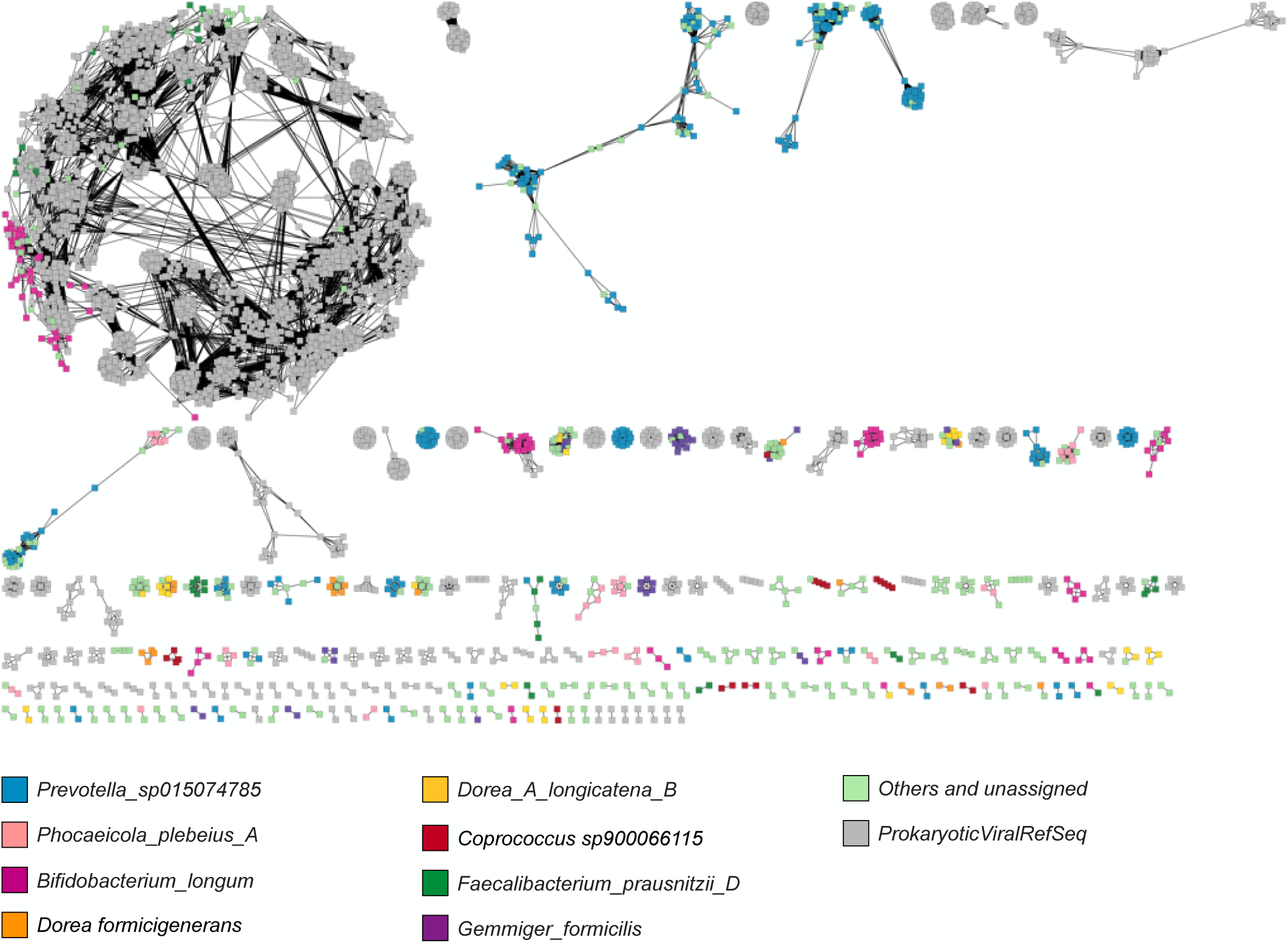
Protein clustering network of prophages detected from bSAGs coupled with Prokaryotic Viral RefSeq Database. Nodes represent individual prophage contigs. Node colors indicate their hosts according to the color legend. Only host bacterial species with more than 30 prophages were denoted.

